# Comparison of Chronic Wasting Disease Detection Methods and Procedures: Implications for Free-Ranging White-Tailed Deer (*Odocoileus Virginianus*) Surveillance and Management

**DOI:** 10.1101/2021.03.03.433751

**Authors:** Marc D. Schwabenlander, Gage R. Rowden, Manci Li, Kelsie LaSharr, Erik C. Hildebrand, Suzanne Stone, Davis M. Seelig, Chris S. Jennelle, Louis Cornicelli, Tiffany M. Wolf, Michelle Carstensen, Peter A. Larsen

## Abstract

Throughout North America, chronic wasting disease (CWD) has emerged as perhaps the greatest threat to wild cervid populations, including white-tailed deer (*Odocoileus virginianus*). White-tailed deer are the most sought after big game species across North America with populations of various subspecies in nearly all Canadian provinces, the contiguous USA, and Mexico. Documented CWD cases have dramatically increased across the white-tailed deer range since the mid-1990s, including in Minnesota. CWD surveillance in free-ranging white-tailed deer and other cervid populations mainly depends upon immunodetection methods (e.g., immunohistochemistry [IHC] and enzyme-linked immunosorbent assay [ELISA]) on medial retropharyngeal lymph nodes and obex. More recent technologies centered on prion protein amplification methods of detection have shown promise as more sensitive and rapid CWD diagnostic tools. Here, we used blinded samples to test the efficacy of real time quaking-induced conversion (RT-QuIC) in comparison to ELISA and IHC for screening tissues, blood, and feces collected in 2019 from white-tailed deer in southeastern Minnesota, where CWD has been routinely detected since 2016. Our results support previous findings that RT-QuIC is a more sensitive tool for CWD detection than current antibody-based methods. Additionally, a CWD testing protocol that includes multiple lymphoid tissues (medial retropharyngeal lymph node, parotid lymph node, and palatine tonsil) per animal may effectively identify a greater number of CWD detections in a white-tailed deer population than a single sample type (i.e., medial retropharyngeal lymph nodes). These results reveal that the variability of CWD pathogenesis, sampling protocol, and testing platform must be considered for the effective detection and management of CWD throughout North America.

## INTRODUCTION

Chronic wasting disease (CWD) is a contagious, 100% fatal neurodegenerative disease affecting deer (*Odocoileus* spp.), moose (*Alces alces*), elk (*Cervus canadensis*), and reindeer (*Rangifer tarandus*). Classified as a transmissible spongiform encephalopathy (TSE), CWD is caused by a misfolded prion protein (PrP^CWD^) which is shed through bodily fluids and can remain infectious in the environment for years (Williams ES, 1980; Prusiner, 1982). Originally detected in Colorado mule deer in 1967, CWD has been detected in additional cervid species and expanded in geographic distribution (Williams and Miller, 2002). As of February 2021, CWD has been found in at least 26 states and three Canadian provinces. The continued expansion of CWD across North America, and recent detections in Scandinavian countries (Mysterud *et al.*, 2020) is changing how cervids are hunted, managed, and consumed. For these reasons, stakeholders tasked with managing CWD must have access to the best diagnostic tools and relevant protocols.

The Minnesota Department of Natural Resources (MNDNR) has surveyed free-ranging white-tailed deer (*Odocoileus virginianus*) for CWD since detection in 2002 in farmed elk and free-ranging white-tailed deer in Minnesota and Wisconsin, respectively. Since then, CWD has been detected in 11 captive cervid facilities and 110 deer in Minnesota (> 90,000 tested), with the disease potentially established endemically in the southeast region, albeit at a low prevalence (La Sharr *et al.*, 2019). Surveillance efforts primarily utilize samples from hunter-harvested animals along with those collected opportunistically from deer found dead in poor body condition, euthanized deer displaying clinical signs of CWD, vehicle collisions, and targeted agency culling.

CWD diagnostic tests can be classified into two categories, first-generation antibody-based diagnostics (e.g., immunohistochemistry [IHC] and enzyme-linked immunosorbent assay [ELISA]) and second-generation prion protein amplification assays (e.g., real time quaking-induced conversion [RT-QuIC] and protein misfolding cyclic amplification [PMCA]). Management agencies have employed ELISA screening of medial retropharyngeal lymph nodes with confirmatory IHC on samples considered suspect by ELISA, as these have traditionally been considered the “gold standard” for CWD (Haley and Richt, 2017). ELISA and IHC, the currently available validated assays for CWD, must be completed at National Animal Health Laboratory Network laboratories. This diagnostic bottleneck has resulted in laboratories being at or beyond testing capacity (Schuler K, Abbott R, Mawdsley J, McGarvey K, January/February 2021). However, advances in CWD diagnostics allow for the implementation of new diagnostic standards and sampling techniques, a critical development as testing pressures and expectations increase (Haley and Richt, 2017; McNulty *et al.*, 2019; Bloodgood *et al.*, 2020; Henderson *et al.*, 2020). Amplification assays have been refined over the past decade and show potential as the next “gold standard” choice of diagnostic tools with increased sensitivity in a high-throughput platform (Haley and Richt, 2017).

We set out to examine the CWD detection capabilities of RT-QuIC test in comparison to ELISA and IHC tests using a population-level sample set of free-ranging deer from southeastern Minnesota. Few studies have investigated the utility of prion amplification assays in comparison to current immunodetection assays on free-ranging cervids (Haley *et al.*, 2014). Importantly, our experimental design included an assessment of RT-QuIC prion detection across multiple tissue types, blood, and feces collected from a subset of sampled individuals. Insights gleaned from this approach will help inform future CWD surveillance efforts and cervid management across North America.

## MATERIALS AND METHODS

### Study area and experimental design

MNDNR contracted with United States Department of Agriculture-Wildlife Services to conduct culling within areas of known CWD-positive deer detections near Preston and Winona, Minnesota; a karst topography region with mixed upland hardwoods, swamp, and agricultural lands (22 January to 29 March 2019; Figure 1). Priority areas were designated spatially by Public Land Survey System sections (1 mi^2^) with a high number of total CWD positive deer, positive female deer (considered to be disease anchors), or areas with high deer densities in close proximity to known positives. Whole, intact carcasses were transported to the Preston DNR Forestry office where MNDNR staff collected tissue samples - medial retropharyngeal lymph nodes (RPLN), submandibular lymph nodes (SLN), parotid lymph nodes (PLN), palatine tonsils (PT), feces, whole blood, and neck muscles (not included in this study). In some cases, significant coagulation of blood or lack of feces precluded collection. All samples were preserved at −20°C in the field.

**Figure 1.**
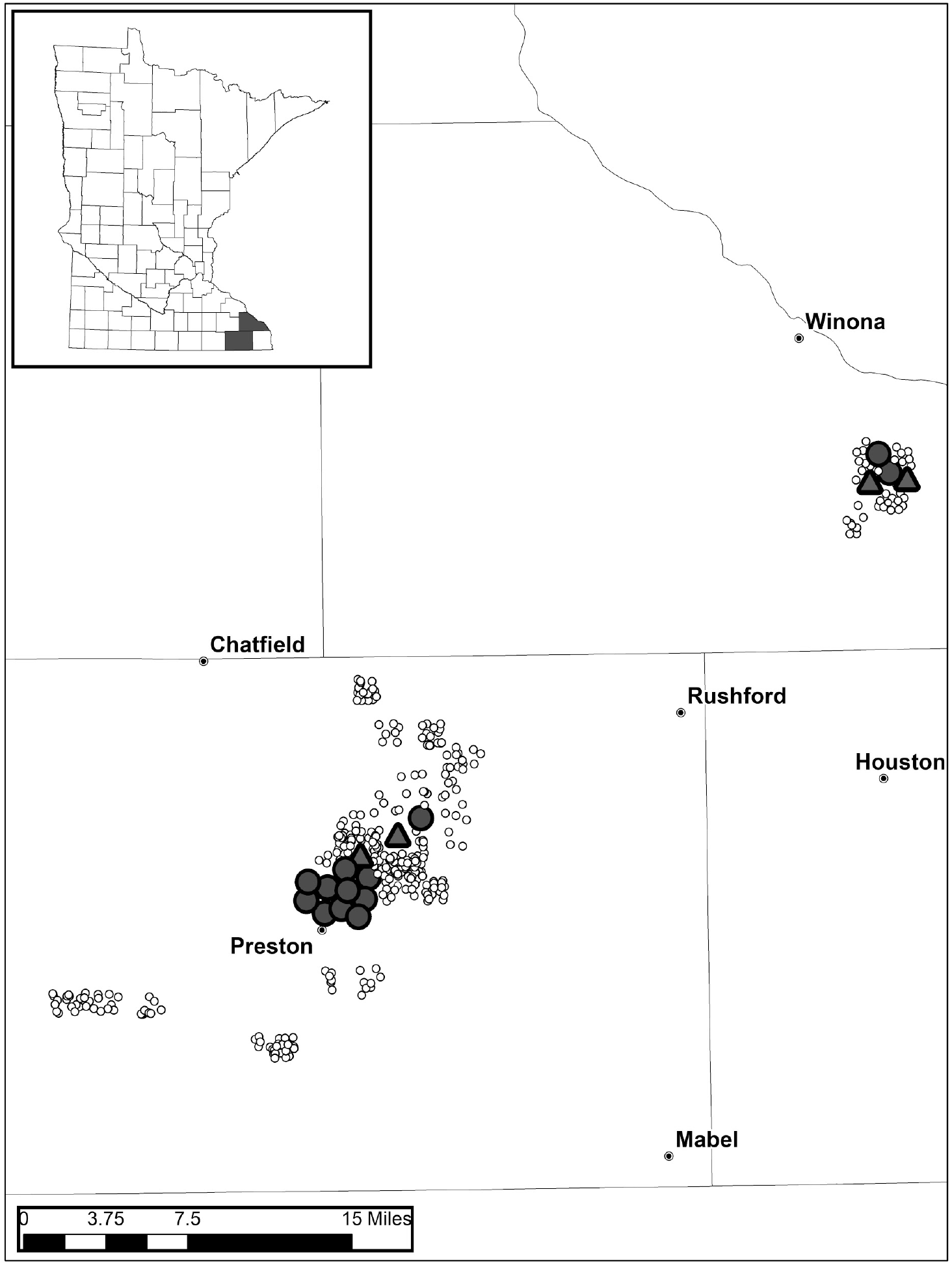
Locations of the 519 white-tailed deer (*Odocoileus virginianus*) collected during the 2019 Minnesota Department of Natural Resources agency culling. Thirteen deer identified as CWD ELISA, IHC, and RT-QuIC (see Results) positive by medial retropharyngeal lymph node samples are indicated by large circles. Four additional deer identified as CWD putative positives by RT-QuIC are indicated by large triangles.

In accordance with surveillance, RPLN samples from all individuals were tested by CWD ELISA at Colorado State University Veterinary Diagnostic Laboratory (CSU VDL) utilizing the Bio-Rad TeSeE Short Assay Protocol (SAP) Combo Kit (BioRad Laboratories Inc., Hercules, CA, USA). For each animal, a pooled homogenate was produced using 3 subsamples from each RPLN. Any suspect-positive RPLN from ELISA was confirmed through IHC of the prion protein as previously described (Hoover *et al.*, 2016).

### RT-QuIC

Following transport from the field, bilateral PLN, SLN, and PT, blood, and feces were preserved at −80°C until RT-QuIC analysis was performed. All RT-QuIC research staff were blinded to the RPLN ELISA/IHC results.

#### Animal/sample identification

RT-QuIC screening began with unilateral sampling and testing of all 519 PLN. Samples that exhibited amyloid seeding activity were tested a second time to confirm the laboratory procedures and both results were reported to the MNDNR. MNDNR staff created a subset for further testing that included all deer that were CWD positive by ELISA/IHC on RPLN, exhibited amyloid seeding activity by RT-QuIC on unilateral PLN, and a randomly chosen set of RPLN ELISA/IHC not detected deer to reach an ELISA/IHC-blinded dataset of 60 animals.

In light of Bloodgood *et al.* (2020), lymphoid tissue sample preparation included bilateral sampling as described below. RT-QuIC testing was performed on bilateral PLN, bilateral SLN, and bilateral PT, as well as the available whole blood and fecal samples for the 60 animal subset. Additionally, the 13 ELISA/IHC CWD positive RPLN were provided by CSU VDL and tested by RT-QuIC. After samples were un-blinded, bilateral PLN, SLN, and PT subsamples of all deer that exhibited amyloid seeding activity on at least one tissue type by RT-QuIC were provided to CSU VDL for ELISA testing. CSU VDL staff were blinded to the RT-QuIC results.

#### Substrate Preparation

Recombinant hamster PrP (HaPrP90-231; provided by NIH Rocky Mountain Laboratory) was cloned into the pET41a(+) expression vector and was expressed in Rosetta™ (DE3) *E. coli* cells (Millipore Sigma, Darmstadt, Germany). Expression and purification of the recombinant substrate was performed following a modified version of the protocol from Orrù *et al.* (2017). Specifically, protein expression was induced using 0.75mM IPTG in place of Overnight Express™ autoinduction (Novagen, Darmstadt, Germany), and cells were cracked with two passes at 16,000 psi on a microfluidizer rather than using a homogenizer.

#### Lymph Tissue Preparation

Lymph tissue (RPLN, PLN, SLN, and PT) dissections were initiated by a cross-sectional cut with sample collection along the cut face to reduce potential cross-contamination that may have originated during field collection. We utilized disposable forceps and scalpels and surface decontamination between samples (1:1.5; 5.25% sodium hypochlorite). Samples were dissected on fresh, disposable benchtop paper. A 10% (w:v) suspension was made by adding 100 mg of tissue to 900 uL of PBS. In the case of bilateral samples, 50 mg of each tissue was dissected and added to one tube. Tissue suspensions were then homogenized using 1.5 mm diameter zirconium oxide beads (Millipore Sigma, Burlington, Massachusetts, USA) and a BeadBug homogenizer (Benchmark Scientific, Sayreville New Jersey, USA), maximum speed for 90 seconds. Homogenized samples were stored at −80°C. Samples were diluted further to 10^−3^ in dilution buffer (0.1% SDS, 1X PBS, N2 supplement), and 2 μL were added to 98 μL of RT-QuIC master mix. The final tissue dilution factor in the reaction was 1:50,000.

#### Blood Preparation

We modified a protocol for blood preparation from Elder *et al.* (2015) and the phosphotungstic acid precipitation was first described by Safar *et al.* (1998). One mL of EDTA whole blood was placed in a tube with 1.5 mm diameter zirconium oxide beads and underwent four cycles of flash freeze-thaw consisting of 3 minutes in dry ice and 3 minutes at 37°C. It was then homogenized using a BeadBug homogenizer at top speed for 2 cycles (a total of 180s). The homogenate was centrifuged at 2,000 rpm for 2 minutes. We incubated 100 μl of supernatant with 7 μl of 4% (w/v) phosphotungstic acid (Sigma-Aldrich, St. Louis, Missouri, USA) in 0.2 M magnesium chloride. The product was then incubated in a ThermoMixer (Eppendorf, Enfield, Connecticut, USA) at 37°C for 1h (1500 rpm) and subsequently centrifuged for 30 minutes at 14,800 rpm. The pellet was resuspended in 20 μL of dilution buffer. We added 2 μl of the 10^−2^–diluted suspension to the RT-QuIC reaction described below.

#### Feces Preparation

We modified a fecal preparation protocol developed by Tennant *et al.* (2020). The fecal pellet was manually homogenized into 10% homogenates using 1X PBS. The solution was centrifuged at 3,000 rpm for 15 minutes at 4°C. We then centrifuged 500 μl of supernatant at 15,000 rpm for 30 minutes. The pellet was resuspended in 100 μl of 1X PBS subsequently incubated with 7 μl of phosphotungstic acid solution as described above. The solution was then centrifuged for 30 minutes at 14,800 rpm. The pellet was resuspended in 10 μl 0.1% SDS in PBS. We added 2 μl of the suspension to the RT-QuIC reaction described below.

#### Assay parameters

Recombinant hamster PrP substrate was filtered at 3,000 x g through a 100 kDa molecular weight cutoff spin column. A master mix was made to the following concentrations: 1X PBS, 500 uM EDTA, 50 uM Thioflavin T, 300 mM NaCl, and 0.1 mg/mL rPrP. All reagents were filter-sterilized through 0.22um PVDF filters. We pipetted 98 uL of the master mix into each well on a black 96-well plate with clear bottoms. After samples were added, the plate was sealed with clear tape. Plates were shaken on a BMG FLUOstar® Omega microplate reader (BMG LABTECH Inc., Cary, North Carolina, USA) at 700 rpm, double orbital for 57 sec and then rested for 83 sec. This shake/rest cycle repeated 21 times, then the fluorescence was recorded. For all tissue types, the temperature was set to 42°C. The whole shake/read cycle would be repeated 58 times for a total of ~46 hr. Readings were recorded with an excitation filter of 450 nm and an emission filter of 480 nm. The gain was set to 1600. The machine performed 21 flashes/well.

#### RT-QuIC Data Analysis

Rate of amyloid formation was determined as the inverse of hours (1/h) for amyloid seeding activity to surpass a threshold of 10 standard deviations above the average baseline readings after 4.5 hr for each plate (Hoover *et al.*, 2016). A minimum of four replicates were performed for each sample. Samples were considered putative positive when at least 50% of the replicates gave a fluorescence signal higher than the threshold cut-off value. P-values were calculated based on rate of amyloid formation versus negative controls using a Mann Whitney U-test as described by Tennant *et al.* (2020) to determine the statistically significant difference in rate of amyloid formation between samples tested and negative controls on the respective plates. Statistical significance was established at 0.05 (α = 0.05) and p-values below 0.05 were considered statistically different. We additionally examined statistical significance for all RT-QuIC data using a maxpoint ratio analysis (Vendramelli *et al.*, 2018; Supplemental File 1). All statistical analyses were performed using GraphPad Prism software (version 9). We report 95% Wilson Score confidence limits for proportions, which is appropriate for small sample size and sample proportions close to 0 or 1 (Brown, Cai and DasGupta, 2001). Confidence limits were estimated using Epitools Epidemiological Calculators (Ausvet, Bruce, ACT, Australia).

## RESULTS

### First generation surveillance: ELISA and IHC

Demographics of the 519 culled deer were as follows: 205 (39%) adult female, 66 (13%) adult male, 36 (7%) yearling female, 53 (10%) yearling male, 82 (16%) fawn female, and 77 (15%) fawn male.

Of the 519 deer, RPLN from 13 deer (0.025; 95% Wilson confidence limits [95% CL] = 0.015 – 0.042) were reported as CWD positive by CSU VDL (Figure 1; Table 1). Demographics of the 13 CWD ELISA/IHC positive deer were as follows: 9 (69%) adult female, 3 (23%) adult male, 0 yearling female, 1 (8%) yearling male, 0 fawn female, and 0 fawn male.

**Table 1.**
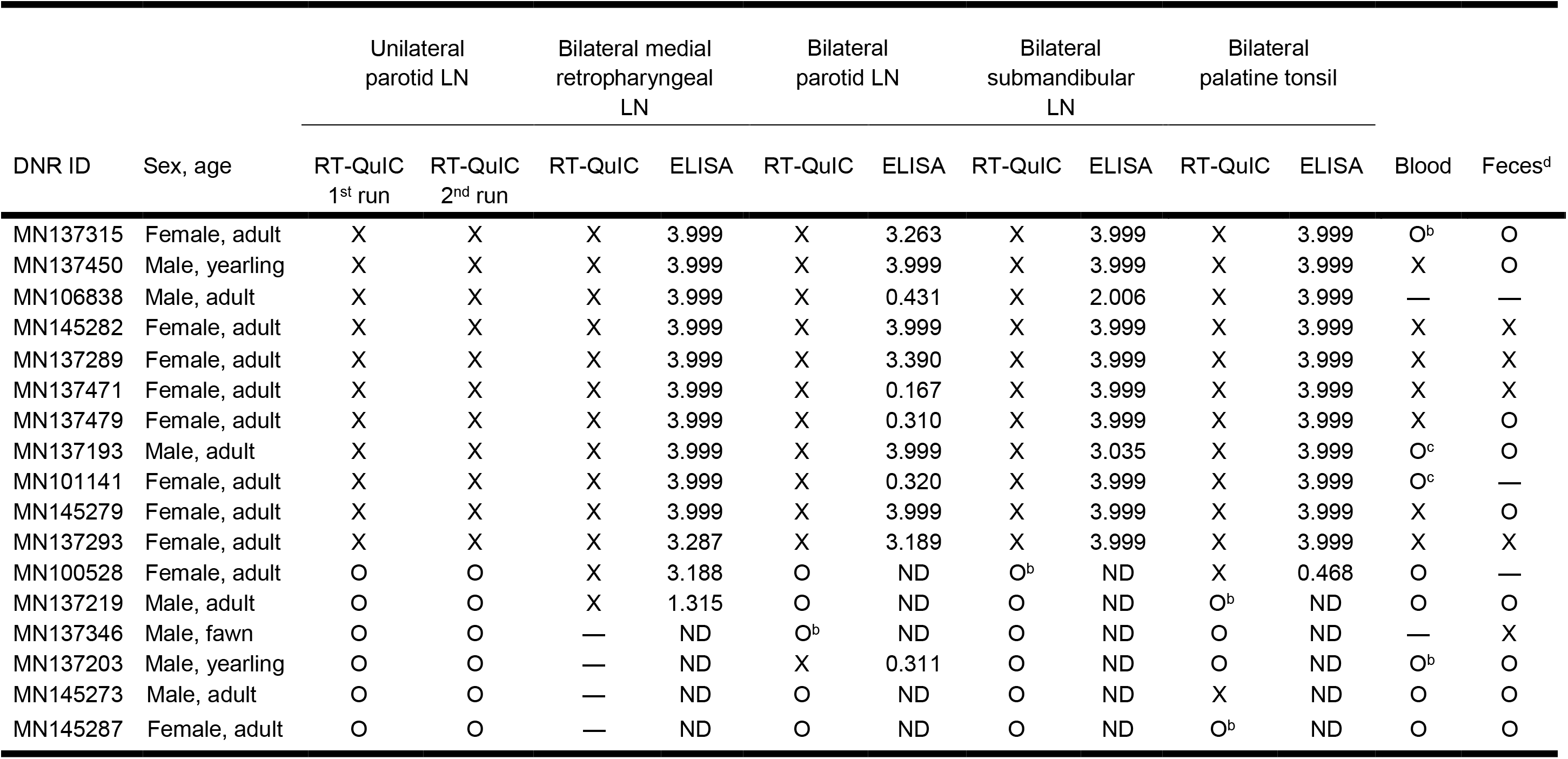
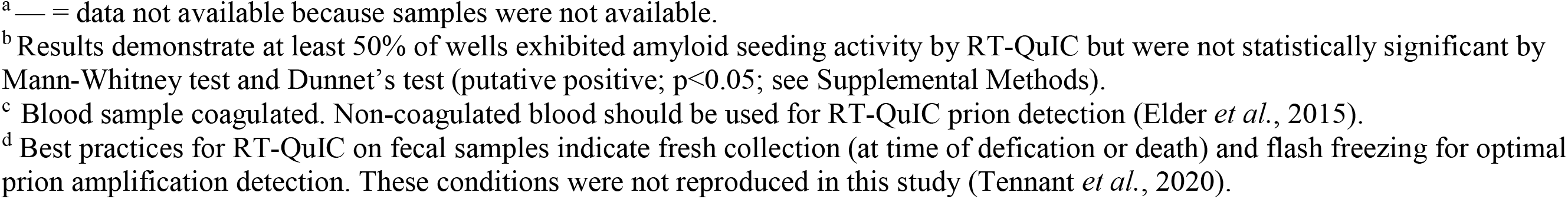
RT-QuIC and ELISA results on animals from the 60 white-tailed deer (*Odocoileus virginianus*) subset that indicated ELISA positivity and/or exhibited amyloid seeding activity by RT-QuIC on at least one sample type.^a^ X=statistically significant amyloid seeding activity determined by Mann-Whitney test and Dunnet’s test (p<0.05; see Supplemental Methods) O=none or not statistically significant amyloid seeding activity determined by Mann-Whitney test and Dunnet’s test (p<0.05; see Supplemental Methods)

| DNR ID | Sex, age | Unilateral parotid LN |  | Bilateral medial retropharyngeal LN |  | Bilateral parotid LN |  | Bilateral submandibular LN |  | Bilateral palatine tonsil |  | Blood | Feces<sup>d</sup> |
| --- | --- | --- | --- | --- | --- | --- | --- | --- | --- | --- | --- | --- | --- |
|  |  | RT-QuIC 1<sup>st</sup> run | RT-QuIC 2<sup>nd</sup> run | RT-QuIC | ELISA | RT-QuIC | ELISA | RT-QuIC | ELISA | RT-QuIC | ELISA |  |  |
| MN137315 | Female, adult | X | X | X | 3.999 | X | 3.263 | X | 3.999 | X | 3.999 | O<sup>b</sup> | O |
| MN137450 | Male, yearling | X | X | X | 3.999 | X | 3.999 | X | 3.999 | X | 3.999 | X | O |
| MN106838 | Male, adult | X | X | X | 3.999 | X | 0.431 | X | 2.006 | X | 3.999 | — | — |
| MN145282 | Female, adult | X | X | X | 3.999 | X | 3.999 | X | 3.999 | X | 3.999 | X | X |
| MN137289 | Female, adult | X | X | X | 3.999 | X | 3.390 | X | 3.999 | X | 3.999 | X | X |
| MN137471 | Female, adult | X | X | X | 3.999 | X | 0.167 | X | 3.999 | X | 3.999 | X | X |
| MN137479 | Female, adult | X | X | X | 3.999 | X | 0.310 | X | 3.999 | X | 3.999 | X | O |
| MN137193 | Male, adult | X | X | X | 3.999 | X | 3.999 | X | 3.035 | X | 3.999 | O<sup>c</sup> | O |
| MN101141 | Female, adult | X | X | X | 3.999 | X | 0.320 | X | 3.999 | X | 3.999 | O<sup>c</sup> | — |
| MN145279 | Female, adult | X | X | X | 3.999 | X | 3.999 | X | 3.999 | X | 3.999 | X | O |
| MN137293 | Female, adult | X | X | X | 3.287 | X | 3.189 | X | 3.999 | X | 3.999 | X | X |
| MN100528 | Female, adult | O | O | X | 3.188 | O | ND | O<sup>b</sup> | ND | X | 0.468 | O | — |
| MN137219 | Male, adult | O | O | X | 1.315 | O | ND | O | ND | O<sup>b</sup> | ND | O | O |
| MN137346 | Male, fawn | O | O | — | ND | O<sup>b</sup> | ND | O | ND | O | ND | — | X |
| MN137203 | Male, yearling | O | O | — | ND | X | 0.311 | O | ND | O | ND | O<sup>b</sup> | O |
| MN145273 | Male, adult | O | O | — | ND | O | ND | O | ND | X | ND | O | O |
| MN145287 | Female, adult | O | O | — | ND | O | ND | O | ND | O<sup>b</sup> | ND | O | O |

<sup>a</sup> — = data not available because samples were not available.
<sup>b</sup> Results demonstrate at least 50% of wells exhibited amyloid seeding activity by RT-QuIC but were not statistically significant by Mann-Whitney test and Dunnet's test (putative positive; $p < 0.05$ ; see Supplemental Methods).
<sup>c</sup> Blood sample coagulated. Non-coagulated blood should be used for RT-QuIC prion detection (Elder *et al.*, 2015).
<sup>d</sup> Best practices for RT-QuIC on fecal samples indicate fresh collection (at time of defecation or death) and flash freezing for optimal prion amplification detection. These conditions were not reproduced in this study (Tennant *et al.*, 2020).

### Second generation surveillance: RT-QuIC

The first objective for RT-QuIC analysis was a blinded screening of the 519 deer using unilateral PLN samples. The first analysis identified 11 (0.021; 95% CL = 0.012 – 0.038) samples exhibiting significant (p<0.05) amyloid seeding activity with repeated results upon retesting (Table 1). We next examined the diagnostic agreement between these 11 animals, animals previously classified as CWD-positive by ELISA/IHC (n=13), and a series of ELISA negative controls (total n=60). One animal that was positive by ELISA/IHC (MN100528) was not included in the 60-deer sample set, but had the same tissue set screened subsequently (following unblinding). Demographics of the 61-deer sample set were as follows: 32 (52%) adult female, 10 (16%) adult male, 5 (8%) yearling female, 4 (7%) yearling male, 6 (10%) fawn female, and 4 (7%) fawn male.

RT-QuIC results indicated 16 (26%) of 61 deer exhibited significant (p<0.05) amyloid seeding activity in at least one sample type. By sample type, we observed amyloid seeding activity in: 12/61 (0.20; 95% CL = 0.12 - 0.31) bilateral PLN, 12/61 (0.20; 95% CL = 0.12 – 0.31) bilateral SLN, 13/61 (0.21; 95% CL = 0.13 – 0.33) bilateral PT, 13/13 (1.0; 95% CL = 0.77 – 1.0) bilateral RPLN, 7/51 (0.14; 95% CL = 0.07 – 0.26) whole blood, and 5/47 (0.11; 95% CL = 0.05 – 0.23) feces (Table 1; Figure 2). There was intra-individual variability in the types of tissues that exhibited amyloid seeding activity for each of the 16 animals (Figures 2,3). We detected amyloid seeding activity beneath levels of significance (p<0.05) in lymphoid and blood samples (n=6) from several of these 16 animals, and considered them putative positive samples (Table 1; Figure 2). Demographics of the 16 deer were as follows: 9 (56%) adult female, 4 (25%) adult male, 0 yearling female, 2 (13%) yearling male, 0 fawn female, and 1 (6%) fawn male. Additionally, amyloid seeding was detected in 2 of 4 replicates (putative positive; not significant) within a palatine tonsil sample from a single animal (MN145287; adult, female) that had no indication of amyloid seeding activity in any other sample type and was ELISA negative across all sample types (Table 1; Figure 2). No other samples from the 61 deer demonstrated amyloid seeding activity.

**Figure 2.**
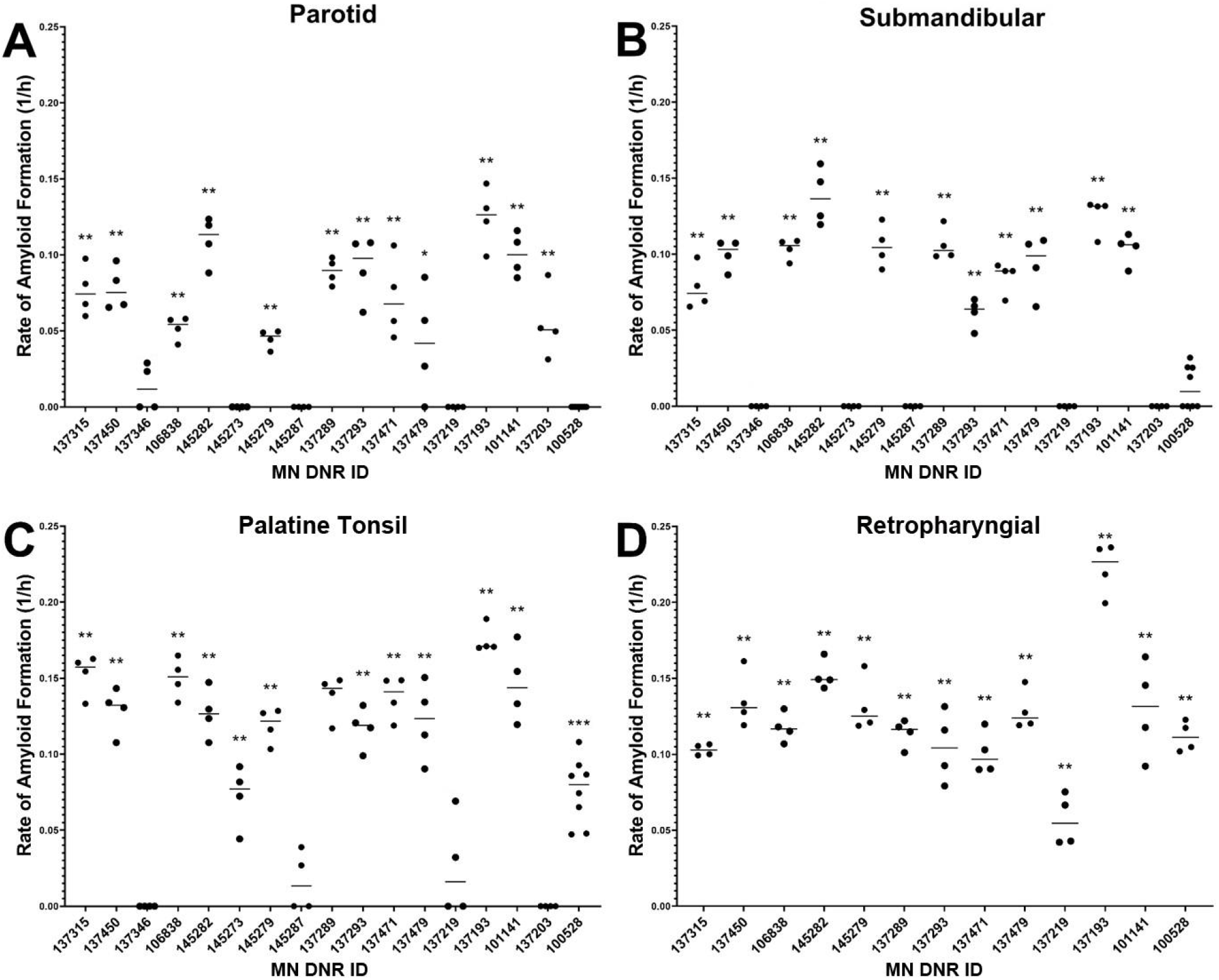
Relative rates of amyloid formation (1/h) of lymphoid tissue samples from the 60 white-tailed deer (*Odocoileus virginianus*) subset that exhibited amyloid seeding activity by RT-QuIC. Samples exhibiting amyloid seeding activity were deemed positive by Mann-Whitney test and Dunnet’s test (***, p < 0.001; **, p < 0.01; *, p < 0.05; see Supplemental Methods).

**Figure 3.**
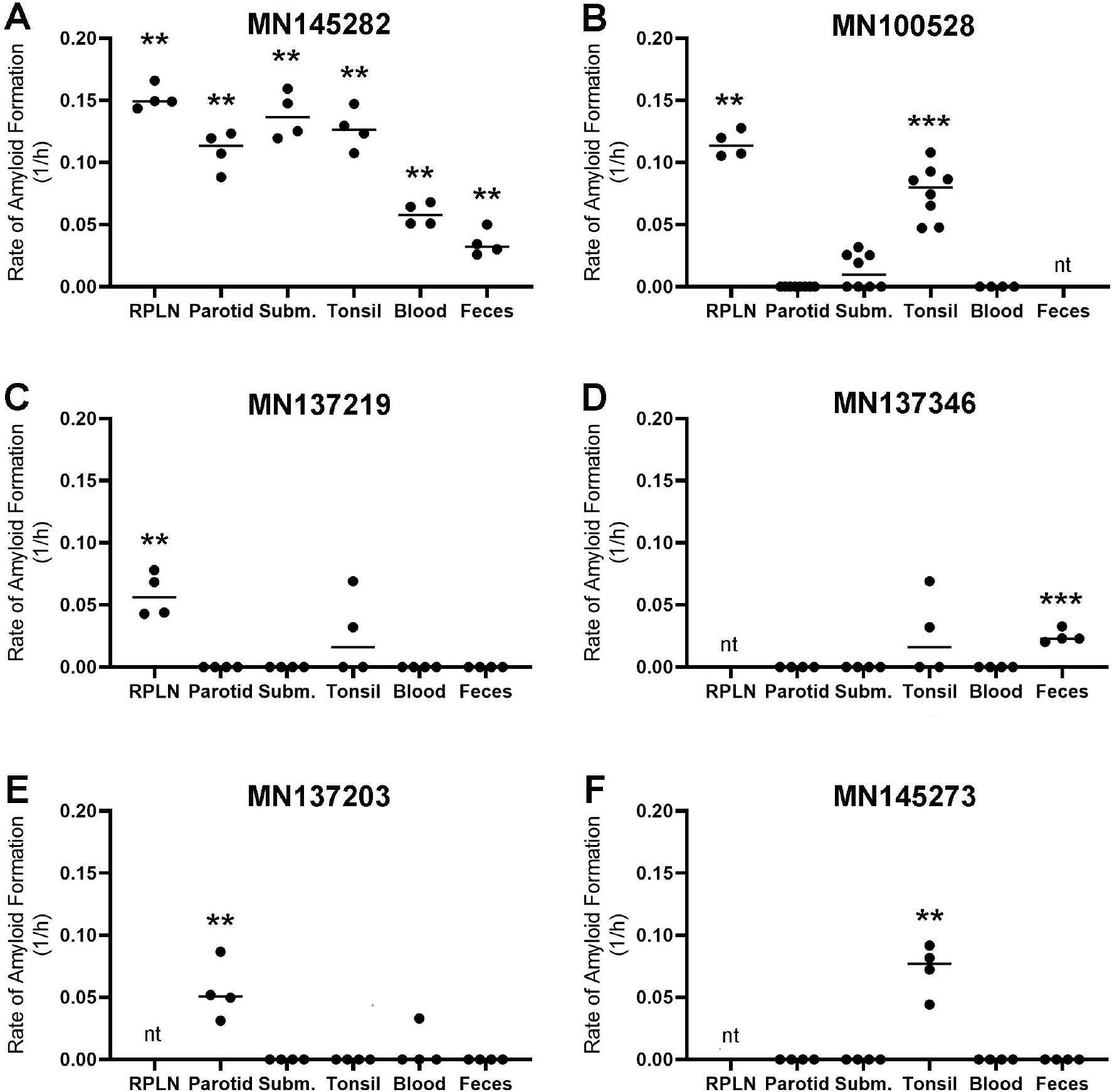
Relative rates of amyloid formation (1/h) of all sample types from six white-tailed deer (*Odocoileus virginianus*) that exhibited amyloid seeding activity by RT-QuIC. Note the variability in amyloid seeding activity across animals. “**A**” demonstrates a highly CWD positive animal across all sample types. “**B**” is likely an animal in earlier disease progression when compared to “**A**”. “**C - F**” are likely animals with very early infection in that only one sample type is statistically significant. Note that “**D - F**” did not have medial retropharyngeal lymph nodes (RPLN) available to test as they were not detected by ELISA and thus discarded at CSU VDL prior to us having access. Samples exhibiting amyloid seeding activity were deemed positive by Mann-Whitney test and Dunnet’s test (***, p < 0.001; **, p < 0.01; *, p < 0.05; see Supplemental Methods). Those samples that were not available for RT-QuIC testing are marked with “nt” (not tested).

### ELISA analysis of animals exhibiting amyloid seeding activity by RT-QuIC

To provide tissue-specific comparison between ELISA and RT-QuIC, PLN, SLN, and PT samples (n=51) from the 17 RT-QuIC positive animals were blind tested by ELISA at CSU VDL. Fifty of the 51 samples demonstrated RT-QuIC and ELISA results agreement (Table 1). MN145273 PT exhibited significant (p<0.05) amyloid seeding activity by RT-QuIC and was not detected by ELISA. Additionally, four samples (MN100528 SLN, MN137219 PT, MN137346 PLN, MN145287 PT) demonstrated amyloid seeding activity that was beneath levels of significance (p<0.05) and were not detected by ELISA. Samples positive on both testing platforms showed a moderate positive correlation of the semi-quantitative measures of prion content (R^2^=0.581; Figure 4). Samples with an ELISA OD value of 3.999 did not bias the correlation as a similar R^2^ was derived from the sample set with those samples removed (data not shown).

**Figure 4.**
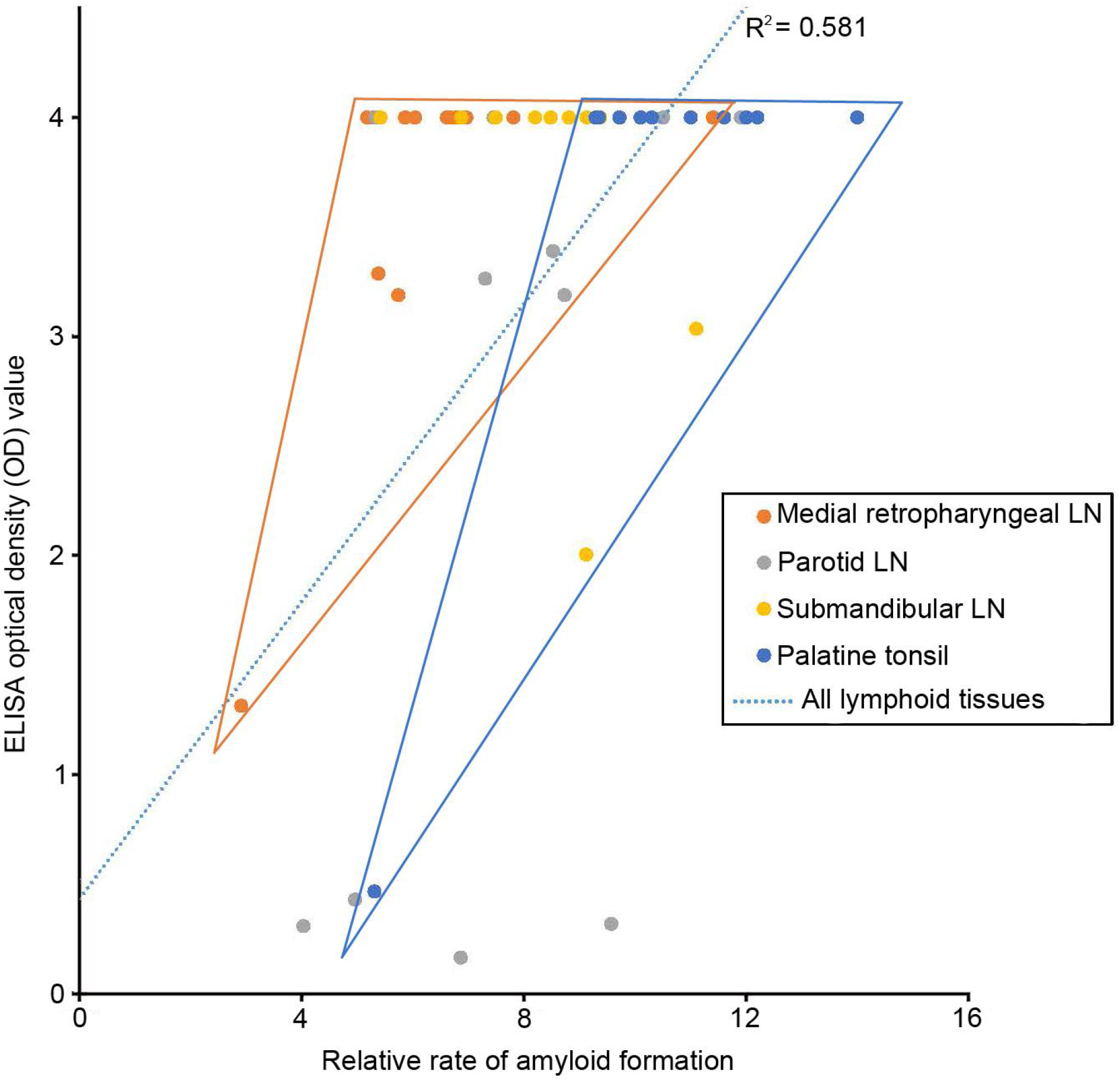
Relationship between the relative rate of amyloid formation and ELISA optical density (OD) for lymphoid tissues examined herein. Depicted samples revealed both amyloid seeding activity by RT-QuIC and had OD values between 0.100 and 3.999. A moderate positive correlation is appreciated across all lymphoid tissue samples (dotted line; Pearson’s correlation coefficient, R^2^ = 0.581). The relationship between medial retropharyngeal lymph node (RPLN) and palatine tonsil (PT) samples is appreciated in that RPLN samples had a generally lower rate of amyloid formation than PT samples.

## DISCUSSION

The goal for diagnostic assays is to optimize the detection of the target so that animals that are diseased or infected are identified as positive (sensitivity) and animals that are not are identified as negative (specificity). An array of factors affect this optimization including, but not limited to, sample type and integrity, sampling method, assay elements, and individual animal (e.g. physiologic or immune state, genetics) and population characteristics (e.g. pathogen prevalence). For CWD, disease detection has primarily been based on ELISA and IHC and the use of postmortem medial retropharyngeal lymph node and obex samples. This approach is effective at identifying animals with end-stage disease, but is limited in detecting low levels of prion burden seen in early stages of disease (Haley and Richt, 2017; Hoover *et al.*, 2017; McNulty *et al.*, 2019). We evaluated the performance of RT-QuIC in comparison to these immunodetection assays with results that support RT-QuIC’s greater potential to detect low levels of CWD prion by utilizing a diverse tissue set that may have value for early detection of the disease.

A growing body of research collectively provides strong evidence that second-generation amplification-based assays have much greater sensitivity in the detection of misfolded PrP than first-generation tests (Haley and Richt, 2017, Soto and Pritzkow, 2018, McNulty *et al.*, 2019). Our results support this conclusion as varied tissue types exhibited amyloid seeding activity with RT-QuIC, but ELISA testing resulted in no detection of PrP^CWD^ in the specific tissue type or within the animal. This led to the discovery of four putative CWD-positive animals by RT-QuIC out of a pool of 48 CWD ELISA not detected animals (RPLN), as well as two ELISA-positive animals (RPLN) with additional putative positive sample types. Collectively, these six animals demonstrated intra-animal variability in CWD positive tissues (ELISA and/or RT-QuIC). Based on relatively low ELISA OD values of the positive tissues in these six animals, and given that RT-QuIC identified tissues that exhibited amyloid seeding activity but were not detected by ELISA, we posit that these six animals were either in the early stages of CWD infection or represent RT-QuIC false-positives. We believe that the former is the case as zero amyloid seeding activity was observed in the 44 “negative” animals (i.e., zero of four replicates) and there was no indication of inter-animal or intra-animal cross-contamination from the sample collection, preparation, or RT-QuIC processes based on the spatial and chronological distribution of CWD detections and the reliability of plate-level controls.

It is problematic to confirm the status of RT-QuIC-positive samples that are negative by first-generation methods as amplification assays approach attogram levels of misfolded prion protein detection, multifold beyond the limits of immunodetection methods (Haley, Richt, *et al.*, 2018). However, similar studies focused on longitudinal analyses of CWD infection support claims that amplification-based assays can effectively detect CWD infections prior to both ELISA and IHC (Haley, Henderson, *et al.*, 2018; Denkers *et al.*, 2020; Haley *et al.*, 2020; Henderson *et al.*, 2020). Previous studies also indicate that the unequal tissue distribution of CWD prion protein in early stages of the disease, coupled with sectioning or sampling technique variability and limitations of immunodetection assays, contributes to the apparent reduced sensitivity of ELISA and IHC (Hoover *et al.*, 2017; Bloodgood *et al.*, 2020). Another complicating factor when comparing first- and second-generation CWD assays, is the potential for variable strains of PrP^CWD^ to be detected by RT-QuIC or PMCA but missed by immunodetection methods due to strain sensitivity to enzymatic digestion. ELISA, IHC, and western-blotting methods utilize antibodies that are unable to distinguish between PrP and PrP^CWD^, thus necessitating Proteinase-K digestion to identify PrP strains resistant to degradation. Recent data indicate that a variety of PrP^CWD^ strains are circulating in cervids, raising the possibility that diagnostic methods which do not utilize enzymatic digestion have greater potential for identifying a broader family of PrP^CWD^ (Duque Velásquez *et al.*, 2015; Osterholm *et al.*, 2019). At the same time, the potential for naturally occurring, non-infectious, conformations of healthy PrP to cause false-positives within amplification-based analyses must be considered.

Sample type also influences CWD test sensitivity, particularly early in infection. RT-QuIC has documented amyloid seeding activity in feces and blood of experimentally-infected deer in the preclinical stages and shortly after inoculation, respectively (M. Elder *et al.*, 2015; Tennant *et al.*, 2020). In both studies, optimal sample conditions - limiting freeze/thaw in feces and heparin preservation in blood - presented the most consistent results. However, these conditions were not reproduced here, which may have contributed to the variable RT-QuIC results from feces and blood. The pathogenesis of CWD provides valuable insight into the prion seeding activity we observed in PT. Tonsillar tissue is one of the first tissues to demonstrate prion immunodeposition, which fits with observed prion traffic through the lymphatic system and is supported in this study (Haley and Richt, 2017; Hoover *et al.*, 2017; Henderson *et al.*, 2020). Palatine tonsil tissue from multiple animals, possibly in early stages of the disease, exhibited amyloid seeding activity by RT-QuIC and had either low OD values or were not detected by ELISA. These results support the value of PT as a tissue of choice for CWD surveillance, particularly when utilizing RT-QuIC. Based on our results, PT demonstrated higher levels of amyloid seeding activity (i.e. “hotter” samples) than did RPLN, yet a majority of both sample types were at the upper limits of detection by ELISA. This leads to the possibility that while RPLN may be good for ELISA-based surveillance, PT may be the ideal tissue type for RT-QuIC-based surveillance. Although the feasibility of identifying PT versus RPLN in field extraction should be considered, further investigation is warranted. We also document the potential for RT-QuIC screening of PLN to identify early stages of CWD infection, and our results supported the continued importance of RPLN for CWD surveillance. Collectively, our RT-QuIC data provide evidence that multi-tissue sampling would provide the best sensitivity for discovering early-infected animals through post-mortem CWD surveillance - likely a combination of PLN, RPLN, and PT, as well as properly collected feces and blood. Fiscal limitations must be accounted for when considering sample type and testing protocols and therefore pooled tissue sample testing may be most appropriate.

Detecting early-infected animals through CWD surveillance is imperative for controlling the disease. This is particularly true for geographic locations where CWD has not been detected. As demonstrated throughout the history of CWD, earlier disease discovery on the landscape leads to earlier implementation of management and more effective control (Miller and Fisher, 2016). The efficacy of tools and protocols for early detection is likely of limited use in core endemic areas where disease is well established, however, implementation of such approaches at the periphery of endemic areas and within areas of incipient CWD expansion events would be critical for detecting and removing newly infected animals, thus limiting the spread of CWD. We posit that the growing number of studies documenting the utility of RT-QuIC for the surveillance of a variety of protein-misfolding and prion diseases, including CWD, collectively demonstrate the method is robust and will greatly aid our understanding and control of CWD (Wilham *et al.*, 2010; Orrú *et al.*, 2015; Caughey *et al.*, 2017; Franceschini *et al.*, 2017; Cooper *et al.*, 2019; Saijo *et al.*, 2019; Henderson *et al.*, 2020; Rossi *et al.*, 2020). In light of the results reported herein, we recommend the implementation of advanced, second-generation, CWD diagnostic assays into existing CWD surveillance, management, and regulatory initiatives.

Future investigation, building on these discoveries, will include optimization of a sampling protocol for CWD amplification assays to increase detection sensitivity, elucidating potential impacts of *PRNP* allele variation on testing for particular CWD strains, and continued analyses of multi-tissue samples secured from CWD positive regions. These efforts, combined with robust statistical validation of new CWD diagnostic tests will help to establish a more complete depiction of the CWD landscape. Our results, utilizing a blinded sample set of free-ranging white-tailed deer from documented CWD hotspots, indicate that RT-QuIC may be a more powerful tool than current methods in identifying CWD positive animals in coordination with a multi-tissue sample collection protocol, particularly for early infections and in CWD-free locations, whether free-ranging or farmed. These findings have direct implications for the effective surveillance and management of CWD across all cervids.

## ACKNOWLEDGEMENTS

We thank NIH Rocky Mountain Labs, especially Byron Caughey, Andrew Hughson, and Christina Orru for training and assistance with the implementation of RT-QuIC. Fred Schendel, Tom Douville, and staff of the University of Minnesota Biotechnology Resource Center provided critical support with respect to large-scale production of recombinant proteins. Kathi Wilson of the Colorado State University Veterinary Diagnostic Laboratory kindly provided assistance with ELISA and IHC testing of samples reported herein. We thank the Minnesota Supercomputing Institute for secure data storage of computational products stemming from our work. Kristen Davenport helped guide our statistical analysis. Lon Hebl graciously provided access to animals housed at the Oxbow Park & Zollman Zoo. This project would not have been possible were it not for the collection of biological samples, and we are grateful to the following persons for their assistance in the field: MNDNR Wildlife staff, Roxanne J. Larsen, Negin Goodarzi, Devender Kumar, Jeremy Schefers, as well as USDA APHIS Wildlife Services staff. Funding for research performed herein was provided by the Minnesota State Legislature through the Minnesota Legislative-Citizen Commission on Minnesota Resources (LCCMR), Minnesota Agricultural Experiment Station Rapid Agricultural Response Fund, University of Minnesota Office of Vice President for Research, Minnesota Department of Natural Resources, and start-up funds awarded to PAL through the Minnesota Agricultural, Research, Education, Extension and Technology Transfer (AGREETT) program.

## Notes

### Competing Interest Statement

The authors have declared no competing interest.

## LITERATURE CITED

Bloodgood, J. et al. (2020) ‘Chronic Wasting Disease Diagnostic Discrepancies: The Importance of Testing Both Medial Retropharyngeal Lymph Nodes’, Journal of wildlife diseases. doi: 10.7589/JWD-D-20-00007.

Brown, L. D., Cai, T. T. and DasGupta, A. (2001) ‘Interval Estimation for a Binomial Proportion’, Statistical science: a review journal of the Institute of Mathematical Statistics, 16(2), pp. 101–117.

Caughey, B. et al. (2017) ‘Amplified Detection of Prions and Other Amyloids by RT-QuIC in Diagnostics and the Evaluation of Therapeutics and Disinfectants’, Progress in molecular biology and translational science, 150, pp. 375–388.

Cooper, S. K. et al. (2019) ‘Detection of CWD in cervids by RT-QuIC assay of third eyelids’, PloS one, 14(8), p. e0221654.

Denkers, N. D. et al. (2020) ‘Very low oral exposure to prions of brain or saliva origin can transmit chronic wasting disease’, PloS one, 15(8), p. e0237410.

Duque Velásquez, C. et al. (2015) ‘Deer Prion Proteins Modulate the Emergence and Adaptation of Chronic Wasting Disease Strains’, Journal of virology, 89(24), pp. 12362–12373.

Elder, A. M. et al. (2015) ‘Immediate and Ongoing Detection of Prions in the Blood of Hamsters and Deer following Oral, Nasal, or Blood Inoculations’, Journal of virology, 89(14), pp. 7421–7424.

Franceschini, A. et al. (2017) ‘High diagnostic value of second generation CSF RT-QuIC across the wide spectrum of CJD prions’, Scientific reports, 7(1), p. 10655.

Haley, N. J. et al. (2014) ‘Detection of chronic wasting disease in the lymph nodes of free-ranging cervids by real-time quaking-induced conversion’, Journal of clinical microbiology, 52(9), pp. 3237–3243.

Haley, N. J., Henderson, D. M., et al. (2018) ‘Chronic wasting disease management in ranched elk using rectal biopsy testing’, Prion, 12(2), pp. 93–108.

Haley, N. J., Richt, J. A., et al. (2018) ‘Design, implementation, and interpretation of amplification studies for prion detection’, Prion, 12(2), pp. 73–82.

Haley, N. J. et al. (2020) ‘Cross-validation of the RT-QuIC assay for the antemortem detection of chronic wasting disease in elk’, Prion, 14(1), pp. 47–55.

Haley, N. J. and Richt, J. A. (2017) ‘Evolution of Diagnostic Tests for Chronic Wasting Disease, a Naturally Occurring Prion Disease of Cervids’, Pathogens, 6(3). doi: 10.3390/pathogens6030035.

Henderson, D. M. et al. (2020) ‘Progression of chronic wasting disease in white-tailed deer analyzed by serial biopsy RT-QuIC and immunohistochemistry’, PloS one, 15(2), p. e0228327.

Hoover, C. E. et al. (2016) ‘Detection and quantification of CWD prions in fixed paraffin embedded tissues by real-time quaking-induced conversion’, Scientific reports, 6, p. 25098.

Hoover, C. E. et al. (2017) ‘Pathways of Prion Spread during Early Chronic Wasting Disease in Deer’, Journal of virology, 91(10). doi: 10.1128/JVI.00077-17.

La Sharr, K. et al. (2019) Surveillance and management of chronic wasting disease in Minnesota. Minnesota Department of Natural Resources.

McNulty, E. et al. (2019) ‘Comparison of conventional, amplification and bio-assay detection methods for a chronic wasting disease inoculum pool’, PloS one, 14(5), p. e0216621.

M. Elder, A. et al. (2015) ‘Longitudinal analysis of blood-borne prion infection’, European Journal of Molecular & Clinical Medicine, 2(4-5), p. 128.

Miller, M. W. and Fisher, J. R. (2016) ‘The First Five (or More) Decades of Chronic Wasting Disease: Lessons for the Five Decades to Come’, in Transactions of the 81st North American Wildlife and Natural Resources Conference; Special Session Three: Science-Based Management Strategies for Fish and Wildlife Diseases. 81st North American Wildlife and Natural Resources Conference, Wildlife Management Institute pp. 110–120.

Mysterud, A. et al. (2020) ‘Policy implications of an expanded chronic wasting disease universe’, The Journal of applied ecology, (1365-2664.13783). doi: 10.1111/1365-2664.13783.

Orrú, C. D. et al. (2015) ‘Rapid and sensitive RT-QuIC detection of human Creutzfeldt-Jakob disease using cerebrospinal fluid’, mBio, 6(1). doi: 10.1128/mBio.02451-14.

Orrù, C. D. et al. (2017) ‘RT-QuIC Assays for Prion Disease Detection and Diagnostics’, Methods in molecular biology, 1658, pp. 185–203.

Osterholm, M. T. et al. (2019) ‘Chronic Wasting Disease in Cervids: Implications for Prion Transmission to Humans and Other Animal Species’, mBio, 10(4). doi: 10.1128/mBio.01091-19.

Prusiner, S. B. (1982) ‘Novel proteinaceous infectious particles cause scrapie’, Science, 216(4542), pp. 136–144.

Rossi, M. et al. (2020) ‘Ultrasensitive RT-QuIC assay with high sensitivity and specificity for Lewy body-associated synucleinopathies’, Acta neuropathologica, 140(1), pp. 49–62.

Safar, J. et al. (1998) ‘Eight prion strains have PrPSc molecules with different conformations’, Nature Medicine, 4(10), pp. 1157–1165.

Saijo, E. et al. (2019) ‘Ultrasensitive RT-QuIC Seed Amplification Assays for Disease-Associated Tau, α-Synuclein, and Prion Aggregates’, Methods in molecular biology, 1873, pp. 19–37.

Schuler K, Abbott R, Mawdsley J, McGarvey K (January/February 2021) ‘Getting results in the fight against CWD’, The Wildlife Professional. Edited by E. Thompson, 15(1), pp. 36–39.

Soto, C. and Pritzkow, S. (2018) ‘Protein misfolding, aggregation, and conformational strains in neurodegenerative diseases’, Nature neuroscience, 21(10), pp. 1332–1340.

Tennant, J. M. et al. (2020) ‘Shedding and stability of CWD prion seeding activity in cervid feces’, PloS one, 15(3), p. e0227094.

Vendramelli, R. et al. (2018) ‘ThermoMixer-Aided Endpoint Quaking-Induced Conversion (EP-QuIC) Permits Faster Sporadic Creutzfeldt-Jakob Disease (sCJD) Identification than Real-Time Quaking-Induced Conversion (RT-QuIC)’, Journal of clinical microbiology, 56(7). doi: 10.1128/JCM.00423-18.

Wilham, J. M. et al. (2010) ‘Rapid end-point quantitation of prion seeding activity with sensitivity comparable to bioassays’, PLoS pathogens, 6(12), p. e1001217.

Williams, E. S. and Miller, M. W. (2002) ‘Chronic wasting disease in deer and elk in North America’, Revue scientifique et technique, 21(2), pp. 305–316.

Williams ES, Y. S. (1980) ‘Chronic wasting disease of captive mule deer: A spongiform encephalopathy’, Journal of wildlife diseases, 16(1), pp. 89–98.

