## Supplemental methods for "Comparison of Chronic Wasting Disease Detection Methods and Procedures: Implications for Free-Ranging White-Tailed Deer (*Odocoileus Virginianus*) Surveillance and Management"

*RT-QuIC Data Analysis - Maxpoint Ratio:* A number of approaches have been implemented to investigate the statistical significance of RT-QuIC data and standardized methods have yet to be broadly defined by the RT-QuIC research community. For this reason, we investigated the utility of the maxpoint ratio as an independent measure to the Mann-Whitney U-test. Significant difference between lymphoid tissue samples and corresponding negative controls was determined based on whether the fluorescence reading from the reaction reaches twice the initial fluorescent intensity. A "maxpoint ratio" was calculated by dividing the maximum fluorescent intensity obtained during the assay by the initial reading (Vendramelli *et al.*, 2018). The rate of amyloid formation was calculated by taking the inverse of the time needed to reach the threshold. For statistical analyses of blood and feces, rate of amyloid formation (1/h) was used to determine whether a sample is statistically different from the corresponding negative control. Threshold was defined as five standard deviations above the average baseline fluorescence. At least four replicates were performed for each sample; p-values were calculated based on the maxpoint ratios or rate of amyloid formation using the Dunnet's test. Significance level was set as 0.05 ($\alpha = 0.05$) and values below 0.05 were considered statistically different.
